# The potent *Toxoplasma gondii* growth inhibitor QQ-437 does not bind to its predicted target, parasite adaptin3*β*, in yeast three-hybrid assays

**DOI:** 10.1101/2022.06.07.495217

**Authors:** Fanny Tran, Jenna E. Foderaro, Nicholas J. Westwood, Gary E. Ward

## Abstract

New drugs are needed to treat infections with *Toxoplasma gondii*, a ubiquitous protozoan parasite that can cause miscarriage, blindness, and life-threatening encephalitis in its human hosts. A novel N-benzoyl-2-hydroxybenzamide named QQ-437 was recently shown to be a potent inhibitor of *T. gondii* growth *in vitro* (EC_50_ = 16nM), and to reduce parasite load in a mouse model of infection. Intriguingly, mutations in the parasite protein TgAP3*β*, which is thought to play an important role in intracellular protein trafficking, confer resistance to QQ-437. In this report, we use yeast three-hybrid analysis to test the hypothesis that QQ-437 inhibits parasite growth via a direct effect on TgAP3*β*. We see no evidence for the ability of QQ-437 to bind directly to TgAP3*β*, suggesting that the mutation in TgAP3*β* that confers resistance does so through an indirect mechanism. Further studies are needed to identify the direct molecular target(s) of this promising class of compounds.

## Introduction

Toxoplasmosis is among the most common parasitic infections of man. Serosurveys show prevalence rates of up to 80% in some parts of the world, and one child out of every 700 born acquires *Toxoplasma* infection *in utero* [1, 2]. Acute infection, while typically subclinical and self-limiting, can be life-threatening in the developing fetus and those who are immunocompromised, and some strains can cause severe disease even in immunocompetent individuals [3]. The toxic and potentially teratogenic effects of currently available drug regimens make management of infection in pregnant woman difficult [4], and while pre-emptive antiparasitic treatment can reduce the risk of toxoplasmic encephalitis in immunocompromised individuals, serious side effects and relapse are common [5-9]. Thus, there is a need for new, better tolerated drugs to prevent or treat clinical toxoplasmosis.

One common approach to antimicrobial drug development is to screen large collections of compounds in cell-based assays for inhibitors of pathogen growth. Once a promising “hit” has been identified, it is advantageous to determine its mechanism of action, *i*.*e*., to identify the target(s) to which it binds. Target identification enables rational, structure-based compound optimization and target-based screening for additional inhibitors that are more potent and/or possess better pharmacokinetic/pharmacodynamic properties; these leads can then be tested in animal models of infection.

Using such an approach, Fomovska and colleagues [10] screened 6,811 compounds with drug-like physicochemical properties [11] for inhibitors of *T. gondii* growth in human fibroblast cells. The screen identified an *N*-benzoyl-2-hydroxybenzamide named MP-IV-1 (Fig. 1A, compound (**1**)), as a potent (EC_50_ = 31nM) parasite growth inhibitor that showed no detectable toxicity to the human fibroblasts. MP-IV-1 (**1**) was also effective at reducing parasitic burden in a mouse model of infection [10]. Subsequent structure-activity relationship analysis identified a closely-related compound, named QQ-437 (**2**) (Fig. 1A, compound (**2**)), as an even more potent, nontoxic inhibitor of parasite growth *in vitro* (EC_50_ = 16nM) and in the mouse model [10]. In an elegant set of experiments, the authors used insertional mutagenesis and selection in QQ-437 (**2**) to generate parasites resistant to the compound. Each of the four resistant clones they isolated had a mutation in the *β* subunit of adaptor protein complex AP-3 (TgAP3*β*) [10]. In other cell types, AP-3 plays an important role in protein sorting at the *trans*-Golgi network and/or endosomes, and in trafficking to the lysosomes [12-14]. It was therefore intriguing that treatment of *T. gondii* with QQ-437 (**2**) led to fragmentation of the lysosome-like compartment in parasites and the development of “empty” dense granule secretory vesicles [10].

**Figure 1:**
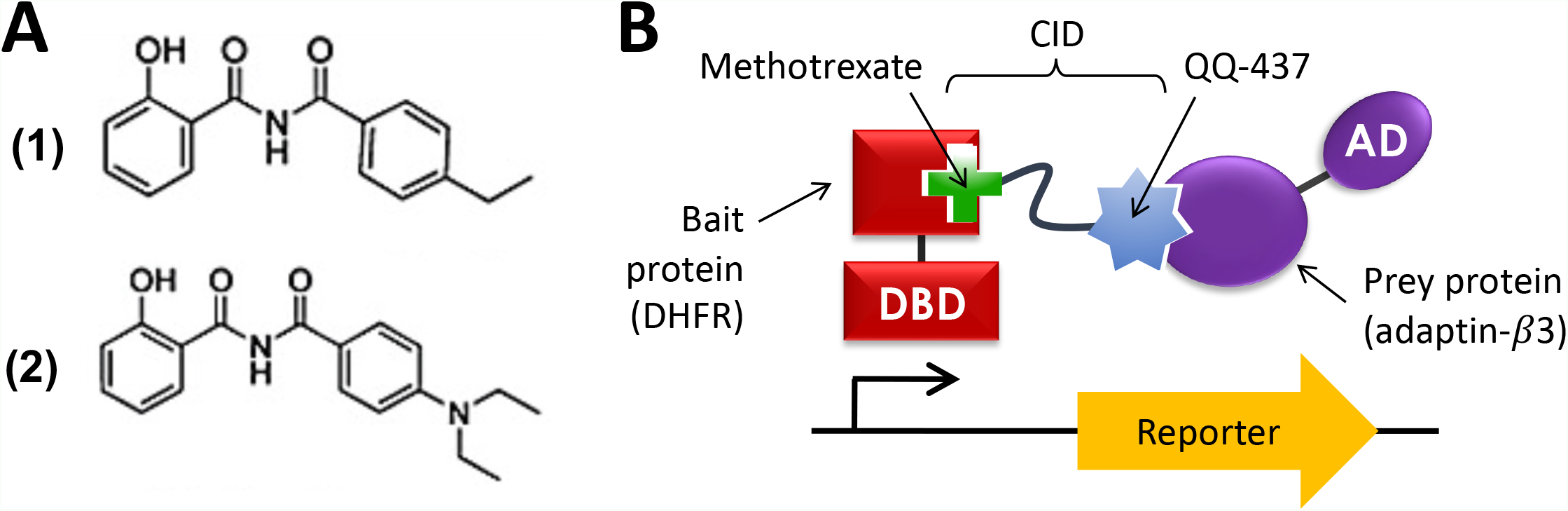
(A) Structures of MP-IV-1 (**1**) and QQ-437 (**2**), identified previously as inhibitors of *T. gondii* growth [10]. (B) Schematic of the yeast 3-hybrid (Y3H) approach to testing whether adaptin-3*β* is a direct target of QQ-437 (**2**). The Gal4 transcription factor is separated into its DNA-binding (DBD) and activation (AD) domains, which are fused to dihydrofolate reductase (DHFR) and adaptin-3*β*, respectively. Upon addition of a chemical inducer of dimerization (CID) composed of methotrexate (which binds to DHFR), a linker, and QQ-437 (**2**). Binding of QQ-437 (**2**) to adaptin-3*β* is expected to reconstitute the transcription factor, thereby activating reporter gene expression. Panel B was modified from [16] and re-used here under the terms of a Creative Commons Attribution License.

Taken together, these data suggest that QQ-437 (**2**) inhibits parasite growth by binding to and inhibiting the function of TgAP3*β*, thereby disrupting critical protein trafficking pathways within the parasite. However, as the authors acknowledged, the data do not exclude the alternative possibility that the mutations in TgAP3*β* confer resistance to the compound through an indirect mechanism, and that the actual target to which the compound binds is something other than TgAP3*β*. It is critical to distinguish between these possibilities before considering any further target-based development/optimization of this scaffold. We therefore set out to test the hypothesis that QQ-437 (**2**) binds directly to TgAP3*β*, using yeast 3-hybrid analysis.

## Methods

### Compound synthesis

All novel analogs of QQ-437 (**2**) and CIDs **5** and **6** were prepared using adapted literature methods [10, 15]. QQ-437 analog **3**: ^1^H NMR (400 MHz, DMSO-*d6*) δ 11.95 (s, 1H), 11.37 (s, 1H), 7.87 (dd, *J* = 8.0, 1.8 Hz, 1H), 7.79 (d, *J* = 9.0 Hz, 1H), 7.43 (t, *J* = 7.4 Hz, 1H), 7.09 – 6.90 (m, 2H), 6.39 (dd, *J* = 9.1, 2.3 Hz, 1H), 6.20 (d, *J* = 2.3 Hz, 1H), 3.96 (s, 3H), 3.43 (q, *J* = 7.0 Hz, 4H), 1.15 (q, *J* = 7.0 Hz, 6H); ^13^C NMR (101 MHz, DMSO-*d6*) δ 164.2, 163.1, 159.9, 157.0, 152.7, 134.4, 134.2, 131.6, 120.0, 119.8, 117.4, 107.9, 104.9, 93.7, 56.0, 44.6, 12.9; HR-MS (ESI) m/z calculated for C_19_H_22_N_2_O_4_Na [M+Na]^+^ 365.1472; found 365.1463.

QQ-437 analog **4**: ^1^H NMR (400 MHz, DMSO-*d6*) δ 12.07 (s, 1H), 11.36 (s, 1H), 7.90 (d, *J* = 8.9 Hz, 1H), 7.77 – 7.68 (m, 2H), 6.77 – 6.68 (m, 2H), 6.64 – 6.50 (m, 2H), 3.80 (s, 3H), 3.43 (q, *J* = 7.0 Hz, 4H), 1.12 (t, *J* = 7.0 Hz, 6H); ^13^C NMR (101 MHz, DMSO-*d6*) δ 164.7, 164.4, 164.3, 159.2, 151.1, 133.2, 130.2, 119.3, 116.4, 111.9, 110.9, 107.3, 101.9, 55.9, 44.3, 12.8; HR-MS (ESI) m/z calculated for C_19_H_22_N_2_O_4_Na [M+Na]^+^: expected 365.1472; found 365.1463.

QQ-437 CID1 **5**: ^1^H NMR (500 MHz, DMSO*-d6*) δ 12.50 (s, 1H), 11.64 (s, 1H), 11.22 (s, 1H), 8.71 (s, 1H), 8.32 (d, *J* = 7.4 Hz, 1H), 8.06 (s, 1H), 7.92 (t, *J* = 5.8 Hz, 1H), 7.77 – 7.67 (m, 4H), 7.48 – 7.39 (m, 1H), 6.98 (d, *J* = 8.4 Hz, 1H), 6.91 (t, *J* = 7.6 Hz, 1H), 6.83 (d, *J* = 8.8 Hz, 2H), 6.48 (d, *J* = 2.2 Hz, 1H), 6.39 (dd, *J* = 9.2, 2.3 Hz, 1H), 5.45 (s, 2H), 4.87 (s, 2H), 4.39 (t, *J* = 5.2 Hz, 2H), 4.32 – 4.24 (m, 1H), 3.68 (t, *J* = 5.2 Hz, 2H), 3.52 – 3.37 (m, 18H), 3.33 (s, 3H), 3.118 – 3.14 (m, 2H), 2.23-2.18 (m, 2H), 2.09 – 2.01 (m, 1H), 1.97 -1.93 (m, 1H), 1.11 (t, *J* = 7.0 Hz, 6H). HR-MS (ESI^-^) m/z calculated for C_51_H_63_N_14_O_12_ [M-H]^-^: 1063.4755; found 1063.4728.

QQ-437 CID2 **6:** ^1^H NMR (700 MHz, DMSO-*d6*) δ 12.50 (br. s, 1H), 11.37 (s, 1H), 8.70 (s, 1H), 8.31 (d, *J* = 7.5 Hz, 1H), 8.22 (s, 1H), 7.94 – 7.88 (m, 1H), 7.76 – 7.65 (m, 4H), 6.85 – 6.77 (m, 2H), 6.76 – 6.68 (m, 2H), 6.68 – 6.64 (m, 2H), 5.19 (s, 2H), 4.85 (s, 2H), 4.58 – 4.51 (m, 2H), 4.32 – 4.26 (m, 1H), 3.86 – 3.78 (m, 2H), 3.54 – 3.30 (m, 22H), 3.25 (s, 3H), 3.21 – 3.14 (m, 2H), 2.24 – 2.16 (m, 2H), 2.08 – 2.00 (m, 1H), 1.96 – 1.88 (m, 1H); 1.42 (t, *J* = 7.0 Hz, 6H); ^13^C NMR (126 MHz, DMSO-*d6*) δ 174.3, 174.1, 172.2, 166.6, 164.7, 164.2, 163.1, 159.0, 158.2, 155.9, 151.1, 149.2, 142.4, 133.2, 131.7, 130.2, 129.4, 125.6, 121.8, 119.2, 112.2, 111.6, 110.9, 107.7, 102.8, 61.8, 55.3, 52.7, 51.5, 49.9, 44.3, 44.2, 39.3, 32.4, 27.0, 12.8. Due to the complexity of the NMR spectrum of (**6**), signals for 2 of the carbon environments were assumed to overlap; HR-MS (ESI) m/z calculated for C_53_H_69_N_14_O_13_ [M+H]^+^: expected 1109.5163; found 1109.5166.

Full analytical data (^1^H, ^13^C, HR-MS) are available on request from NJW.

### Y3H analysis

Y3H assays were performed as previously described [16], with the following modifications. The coding sequence of TgAP3*β* was cloned into pGADT7 (Clontech Laboratories, Mountain View, CA) and co-transformed into competent AH109 yeast cells with a plasmid encoding *E. coli* dihydrofolate reductase fused to the Gal4 DNA-binding domain.

The transformants were plated onto yeast drop out plates (SC +Glu –trp –leu) and incubated at 30°C for 48 hours. A single colony of the resulting transformants was grown overnight in 3 ml of synthetic complete liquid media (SC +Glu –trp –leu). Cells were sub-cultured and grown to log phase in 4ml of SC +Glu –trp –leu (determined by OD600) and diluted to 2×105 cells/ml. Each test well of a 96-well plate (BD Falcon) containing 125 μl SC +Glu – trp –leu –his was inoculated with 25 μl of cell suspension. CIDs (1.5 μl of 2.5 mM) and 3-AT (1.5 μl of 50 mM) were added to test wells. As a negative control, 1.5 μl DMSO and 1.5 μl 3-AT were added to wells containing 150 μl of media alone. Plates were parafilmed and incubated with shaking (200 rpm) at 30°C. The OD600 for each well was recorded daily after mixing using a Synergy 2 microplate reader (BioTek, Winooski, VT).

## Results and Discussion

When a potential target of a small molecule of interest has been identified, yeast three-hybrid (Y3H) analysis provides a powerful approach to confirm direct binding of the small molecule to the putative target [16, 17]. Y3H analysis is a modified version of the more familiar yeast two-hybrid system [15, 17-20]. In Y3H, bait and prey proteins (each fused to one part of a transcription factor) interact indirectly, through a bridging molecule called a Chemical Inducer of Dimerization (CID; Fig. 1B). The bivalent CID consists of two small molecules connected by a flexible linker: one small molecule binds to the bait protein and the other to the prey. The formation of a ternary complex consisting of bait, CID and prey leads to reconstitution of the transcription factor and activation of one or more reporter genes (Fig. 1B).

We used Y3H to test the hypothesis that QQ-437 (**2**) binds to *T. gondii* adaptin-*β*3. In the Y3H system used here, the prey protein consists of dihydrofolate reductase (DHFR) fused to the DNA-binding domain of the GAL4 transcription factor. DHFR binds to methotrexate at one end of the CID and the other end of the CID displays QQ-437 (**2**). *T. gondii* adaptin-3*β* fused to the activation domain of GAL4 is then expressed in these cells, serving as the prey. If addition of the QQ-437-containing CID leads to yeast cell growth via the activation of the required yeast reporter gene, binding of QQ-437 (**2**) to adaptin-3*β* will be confirmed (Fig. 1A).

To synthesize the CID(s) for use in these studies we undertook a structure-activity relationship (SAR) study to determine where an OMe group could be incorporated into QQ-437 (**2**) for linker attachment without losing biological activity. Resynthesized QQ-437 (**2**) inhibited parasite growth with an IC_50_ of 4nM. Of the six possible novel mono-methoxy analogs, all of which were prepared, one (Fig. 2A, compound (**3**)) showed a nearly 50-fold increase in potency in the parasite growth assay (EC_50_ = 90 pM) compared to QQ-437 (**2**). A second analog (**4**) showed only a minor decrease in potency compared to QQ-437 (**2**) (EC_50_ = 16 nM). We therefore synthesized two CIDs, QQ-437 CID1 (**5**) and QQ-437 CID2 (**6**), containing methotrexate linked to (**3**) or (**4**), respectively (Fig. 2B).

**Figure 2:**
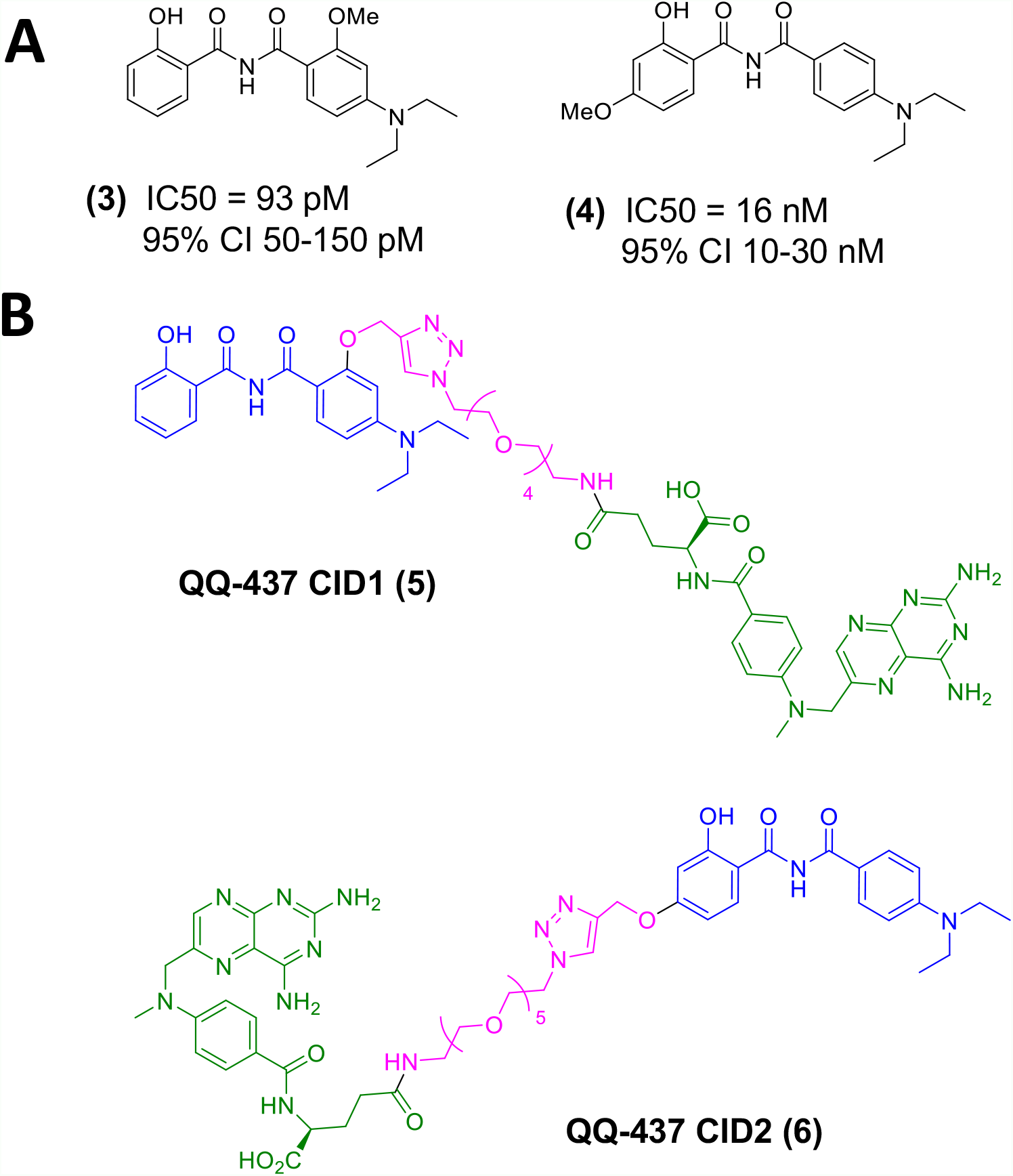
Structure of the CIDs used in this study. (A) Structure and biological activity of the two QQ-437 analogs on which the syntheses of the two CIDs were based. (B) Structures of QQ-437 CID1 (**5**) and QQ-437 CID2 (**6**). QQ-437 component is highlighted in blue, linker in pink, and methotrexate is shown in green (*c*.*f*. Figure 1B).

We then cloned TgAP3*β* into the Y3H vector pGADT7 and used CIDs (**5**) and (**6**) to test for a direct interaction between QQ-437 (**2**) and TgAP3*β*. As a positive control, we cloned the parasite protein TgBradin into the same Y3H vector and tested for growth in the presence of MTX-Cmpd2.1, a CID previously shown by Y3H to bind to TgBradin [16]. The positive control showed robust yeast growth at both 48 and 72 hr compared to the DMSO vehicle alone (Fig. 3A and B). In contrast, no growth above background was seen using equivalent amounts of either QQ-437 CID1 (**5**) (Fig. 3A) or QQ-437 CID2 (**6**) (Fig. 3B) in yeast expressing TgAP3*β*, strongly suggesting a lack of direct interaction between QQ-437 (**2**) and TgAP3*β*.

**Figure 3:**
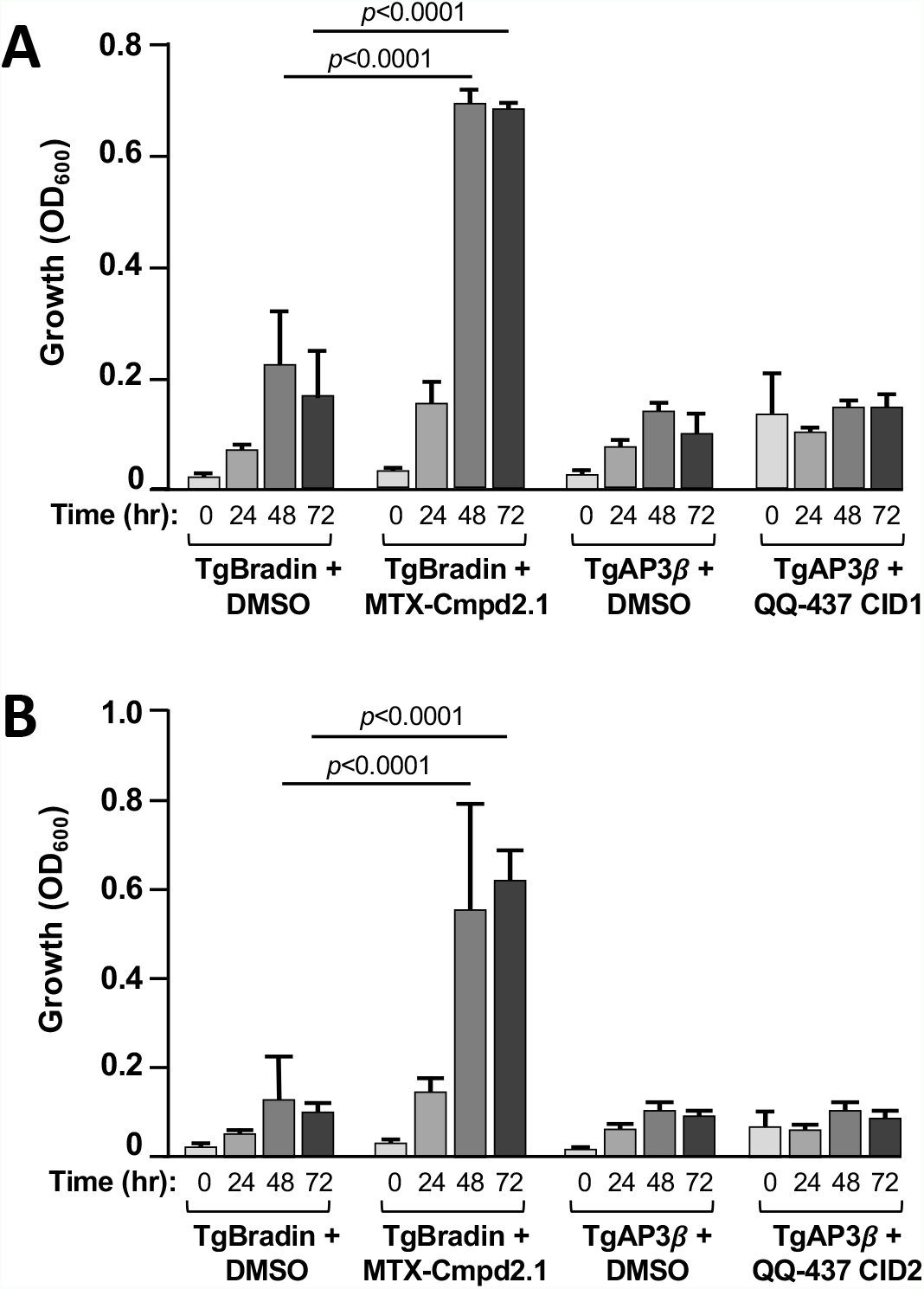
Targeted yeast three-hybrid suggests QQ-437 (**2**) does not interact with TgAP3*β*. Growth of yeast is a reflection of ternary complex formation and reporter activation. Yeast expressing the positive control plasmid, pGAD-TgBRADIN, grow more robustly in the presence of the CID MTX-Cmpd2.1 than in DMSO, demonstrating an interaction between TgBradin and MTX-Cmpd2.1, as shown previously [16]. In contrast, yeast expressing pGAD-AP3B grow to the same extent in (A) QQ-437 CID1 (**5**), (B) QQ-437 CID2 (**6**), and DMSO (A, B), indicating no binding of either CID to TgAP3*β*. Bars represent mean +/-standard deviation; n=3. Data were analyzed using a Student’s paired t-test (\*\*\*\**p*≤0.0001).

The generation of parasites resistant to a compound of interest and identification of mutations linked to the resistance phenotype is a proven and powerful approach to target identification (*e*.*g*., [21-23]). However, a gene so identified may encode the direct target of the compound, another component of the pathway in which the target functions, or a protein that functions in resistance independent of the function of the target itself (*e*.*g*., a transporter protein that extrudes the compound from the parasite). Our data suggest that TgAP3*β* is not in fact a direct target for QQ-437.

## Limitations

One caveat to this conclusion is that attachment of the linker/DHFR to QQ-437 (**2**) may have abrogated its TgAP3*β*-binding activity. While we cannot rule out this possibility, we generated two independent QQ-437-based CIDs, with the linker attached at positions that our SAR suggested could tolerate substitution, and neither showed detectable binding to TgAP3*β* by Y3H. Furthermore, this class of compounds also inhibits the growth of the related parasite, *Plasmodium falciparum* [24], yet the *β* subunit of AP3 is either absent or highly degenerate in *Plasmodium spp*. [25]. Thus, while we cannot rule out TgAP3*β* as a direct target, the results presented here cast doubt on the hypothesis that QQ-437 inhibits parasite growth through a direct effect on TgAP3*β*, and they call for further experiments to definitively identify the target(s) of this promising class of parasite growth inhibitors.

## Declarations

### Ethics approval/consent to participate

N/A

### Availability of data and material

Yeast strains available on request from GEW. Full analytical data (^1^H, ^13^C, HR-MS) on the synthesized compounds available on request from NJW.

## Acknowledgements

We thank Virginia Cornish for providing yeast strains.

## Consent for publication

N/A

## Author’s contributions

Conception and design of the experiments – FT, JF, NJW, GEW; Data generation – FT, JF; Analysis and interpretation of data – FT, JF, NJW, GEW; Drafting the manuscript –NJW, GEW; Editing the manuscript – FT, JF.

## References

1. Pappas G, Roussos N, Falagas ME. Toxoplasmosis snapshots: global status of Toxoplasma gondii seroprevalence and implications for pregnancy and congenital toxoplasmosis. Int J Parasitol. 2009;39(12):1385–94. Epub 2009/05/13. doi: 10.1016/j.ijpara.2009.04.003. PubMed PMID: 19433092.

2. Torgerson PR, Mastroiacovo P. The global burden of congenital toxoplasmosis: a systematic review. Bull World Health Organ. 2013;91(7):501–8. Epub 2013/07/05. doi: 10.2471/BLT.12.111732. PubMed PMID: 23825877; PubMed Central PMCID: PMC3699792.

3. Dubey JP. Outbreaks of clinical toxoplasmosis in humans: five decades of personal experience, perspectives and lessons learned. Parasites & vectors. 2021;14(1):263. Epub 2021/05/21. doi: 10.1186/s13071-021-04769-4. PubMed PMID: 34011387.

4. Remington JS, McLeod R, Desmonts G. Toxoplasmosis. In: Remington JS, Klein JO, editors. Infectious Diseases of the Fetus and the Newborn Infant. 4th ed. Philadelphia: W. B. Saunders Company; 1995. p. 140–267.

5. Ben-Harari RR, Goodwin E, Casoy J. Adverse Event Profile of Pyrimethamine-Based Therapy in Toxoplasmosis: A Systematic Review. Drugs R D. 2017;17(4):523–44. doi: 10.1007/s40268-017-0206-8. PubMed PMID: 28879584; PubMed Central PMCID: PMCPMC5694419.

6. Choquet-Kastylevsky G, Vial T, Descotes J. Allergic adverse reactions to sulfonamides. Curr Allergy Asthma Rep. 2002;2(1):16–25. PubMed PMID: 11895621.

7. McLeod R, Khan AR, Noble GA, Latkany P, Jalbrzikowski J, Boyer K, et al. Severe sulfadiazine hypersensitivity in a child with reactivated congenital toxoplasmic chorioretinitis. Pediatr Infect Dis J. 2006;25(3):270–2. doi: 10.1097/01.inf.0000202070.59190.9a. PubMed PMID: 16511396.

8. McPhillie M, Zhou Y, El Bissati K, Dubey J, Lorenzi H, Capper M, et al. New paradigms for understanding and step changes in treating active and chronic, persistent apicomplexan infections. Scientific reports. 2016;6:29179. doi: 10.1038/srep29179. PubMed PMID: 27412848; PubMed Central PMCID: PMCPMC4944145.

9. Porter SB, Sande MA. Toxoplasmosis of the central nervous system in the acquired immunodeficiency syndrome. N Engl J Med. 1992;327(23):1643–8. Epub 1992/12/03. doi: 10.1056/NEJM199212033272306. PubMed PMID: 1359410.

10. Fomovska A, Huang Q, El Bissati K, Mui EJ, Witola WH, Cheng G, et al. Novel N-benzoyl-2-hydroxybenzamide disrupts unique parasite secretory pathway. Antimicrob Agents Chemother. 2012;56(5):2666–82. Epub 2012/02/23. doi: 10.1128/AAC.06450-11. PubMed PMID: 22354304; PubMed Central PMCID: PMC3346615.

11. Lipinski CA, Lombardo F, Dominy BW, Feeney PJ. Experimental and computational approaches to estimate solubility and permeability in drug discovery and development settings. Adv Drug Deliv Rev. 2001;46(1-3):3–26. PubMed PMID: 11259830.

12. Cowles CR, Odorizzi G, Payne GS, Emr SD. The AP-3 adaptor complex is essential for cargo-selective transport to the yeast vacuole. Cell. 1997;91(1):109–18. Epub 1997/10/23. doi: 10.1016/s0092-8674(01)80013-1. PubMed PMID: 9335339.

13. Feraru E, Paciorek T, Feraru MI, Zwiewka M, De Groodt R, De Rycke R, et al. The AP-3 β adaptin mediates the biogenesis and function of lytic vacuoles in Arabidopsis. Plant Cell. 2010;22(8):2812–24. Epub 2010/08/24. doi: 10.1105/tpc.110.075424. PubMed PMID: 20729380; PubMed Central PMCID: PMCPMC2947184.

14. Zwiewka M, Feraru E, Möller B, Hwang I, Feraru MI, Kleine-Vehn J, et al. The AP-3 adaptor complex is required for vacuolar function in Arabidopsis. Cell Res. 2011;21(12):1711–22. Epub 2011/06/15. doi: 10.1038/cr.2011.99. PubMed PMID: 21670741; PubMed Central PMCID: PMCPMC3357998.

15. Tran F, Odell AV, Ward GE, Westwood NJ. A modular approach to triazole-containing chemical inducers of dimerisation for yeast three-hybrid screening. Molecules. 2013;18(9):11639–57. doi: 10.3390/molecules180911639. PubMed Central PMCID: PMC4031444.

16. Odell AV, Tran F, Foderaro JE, Poupart S, Pathak R, Westwood NJ, et al. Yeast Three-Hybrid Screen Identifies TgBRADIN/GRA24 as a Negative Regulator of Toxoplasma gondii Bradyzoite Differentiation. PLOS One. 2015;10(3):e0120331. Epub 2015/03/20. doi: 10.1371/journal.pone.0120331. PubMed PMID: 25789621; PubMed Central PMCID: PMC4366382.

17. Becker F, Murthi K, Smith C, Come J, Costa-Roldan N, Kaufmann C, et al. A three-hybrid approach to scanning the proteome for targets of small molecule kinase inhibitors. Chem Biol. 2004;11(2):211–23. PubMed PMID: 15123283.

18. Baker K, Sengupta D, Salazar-Jimenez G, Cornish VW. An optimized dexamethasone-methotrexate yeast 3-hybrid system for high-throughput screening of small molecule-protein interactions. Anal Biochem. 2003;315(1):134–7. PubMed PMID: 12672422.

19. Chidley C, Haruki H, Pedersen MG, Muller E, Johnsson K. A yeast-based screen reveals that sulfasalazine inhibits tetrahydrobiopterin biosynthesis. Nat Chem Biol. 2011;7(6):375–83. PubMed PMID: 21499265.

20. Licitra EJ, Liu JO. A three-hybrid system for detecting small ligand-protein receptor interactions. Proc Natl Acad Sci U S A. 1996;93(23):12817–21.

21. Istvan ES, Dharia NV, Bopp SE, Gluzman I, Winzeler EA, Goldberg DE. Validation of isoleucine utilization targets in Plasmodium falciparum. Proc Natl Acad Sci U S A. 2011;108(4):1627–32. PubMed PMID: 21205898.

22. Rosenberg A, Luth MR, Winzeler EA, Behnke M, Sibley LD. Evolution of resistance in vitro reveals mechanisms of artemisinin activity in Toxoplasma gondii. Proc Natl Acad Sci U S A. 2019. Epub 2019/12/07. doi: 10.1073/pnas.1914732116. PubMed PMID: 31806760.

23. Wu W, Herrera Z, Ebert D, Baska K, Cho SH, DeRisi JL, et al. A chemical rescue screen identifies a Plasmodium falciparum apicoplast inhibitor targeting MEP isoprenoid precursor biosynthesis. Antimicrob Agents Chemother. 2015;59(1):356–64. Epub 2014/11/05. doi: 10.1128/aac.03342-14. PubMed PMID: 25367906; PubMed Central PMCID: PMCPMC4291372.

24. Stec J, Huang Q, Pieroni M, Kaiser M, Fomovska A, Mui E, et al. Synthesis, biological evaluation, and structure-activity relationships of N-benzoyl-2-hydroxybenzamides as agents active against P. falciparum (K1 strain), Trypanosomes, and Leishmania. J Med Chem. 2012;55(7):3088–100. Epub 2012/02/23. doi: 10.1021/jm2015183. PubMed PMID: 22352841; PubMed Central PMCID: PMCPMC3330251.

25. Nevin WD, Dacks JB. Repeated secondary loss of adaptin complex genes in the Apicomplexa. Parasitol Int. 2009;58(1):86–94. Epub 2009/01/17. doi: 10.1016/j.parint.2008.12.002. PubMed PMID: 19146987.

